# Distinct mitochondrial phenotypes align with visual and semantic representations across human cortex

**DOI:** 10.64898/2026.08.07.743627

**Authors:** Zitong Lu, Yuxin Wang

**Affiliations:** McGovern Institute for Brain Research at Massachusetts Institute of Technology, MA, USA; Pritzker School of Molecular Engineering at University of Chicago, IL, USA

## Abstract

Cortical regions differ in the information they represent, but whether this functional specialization is accompanied by corresponding differences in cellular and molecular organization remains unknown. Here, we combined precision 7T fMRI, image-to-brain encoding models, a recently developed human mitochondrial atlas, and cortical transcriptomics to test whether visual and semantic representations occupy distinct molecular environments. Across two independent combinations of visual and semantic feature models, variance uniquely attributable to visual information was negatively associated with both mitochondrial density and mitochondria-specific respiratory capacity. In contrast, variance uniquely attributable to semantic information was positively associated with mitochondrial density but showed no reliable association with respiratory capacity. Genome-wide transcriptomic analyses revealed convergent differences in mitochondrial and broader cellular programs associated with the two representational axes. These findings establish a cross-scale correspondence between the information represented by human cortex and its regional energetic and molecular architecture.

## Introduction

Neural computation imposes substantial metabolic demands, yet the biological organization that sustains cortical computation is typically studied separately from the information represented by neural populations. Human neuroimaging has revealed systematic transitions from sensory to increasingly abstract representations across cortex, whereas molecular and cellular studies have demonstrated pronounced regional variation in mitochondrial abundance, respiratory properties, gene expression, and cell composition. However, the energetic organization of the human brain has only been investigated in relation to anatomical variation, disorders, and macroscale network organization (1–6), leaving unresolved how it relates to the specific information represented during human cognition. To our knowledge, no study has jointly tested whether the cortical strength of distinct information representations is spatially aligned with regional mitochondrial phenotypes and associated with coordinated cortical transcriptional programs.

Mitochondria provide a compelling focus because they are central to cellular bioenergetics and generate much of the ATP required to sustain neuronal signaling (7). Two complementary postmortem resources make such a multiscale test possible. A recent mitochondrial atlas (8) mapped, among other properties, mitochondrial density (MitoD), a measure of regional mitochondrial abundance, and mitochondrial respiratory capacity (MRC), a measure of mitochondrial specialization for OXPHOS energy transformation on a per-mitochondrion basis. In parallel, the Allen Human Brain Atlas (AHBA) (9) provides spatially annotated, genome-wide gene-expression measurements across human cortex, enabling the identification of mitochondrial and broader cellular programs associated with the same representational axes. Here, using 7T fMRI during natural-scene viewing (10) and two image-to-brain encoding frameworks, we separated cortical variance uniquely attributable to visual and semantic features and tested its correspondence with both regional mitochondrial phenotypes and cortical transcriptional organization.

## Results

### Visual and semantic information define robust cortical representational axes

We first derived participant-specific cortical maps of information representation during natural-scene viewing. We separated image-evoked cortical responses into variance uniquely captured by visual features and variance uniquely captured by semantic descriptions. Four participants from the Natural Scenes Dataset each viewed approximately 10,000 images three times. After averaging repeats, we trained encoding models on participant-specific images using visual and semantic embeddings from deep artificial neural networks (ANNs) and evaluated on the 1,000 images shared by all participants. Visual-only, semantic-only and joint predictions were combined through nested cross-validated ridge regression and convex stacking; variance partitioning then yielded unique visual and unique semantic maps (Figure 1A).

**Figure 1.**
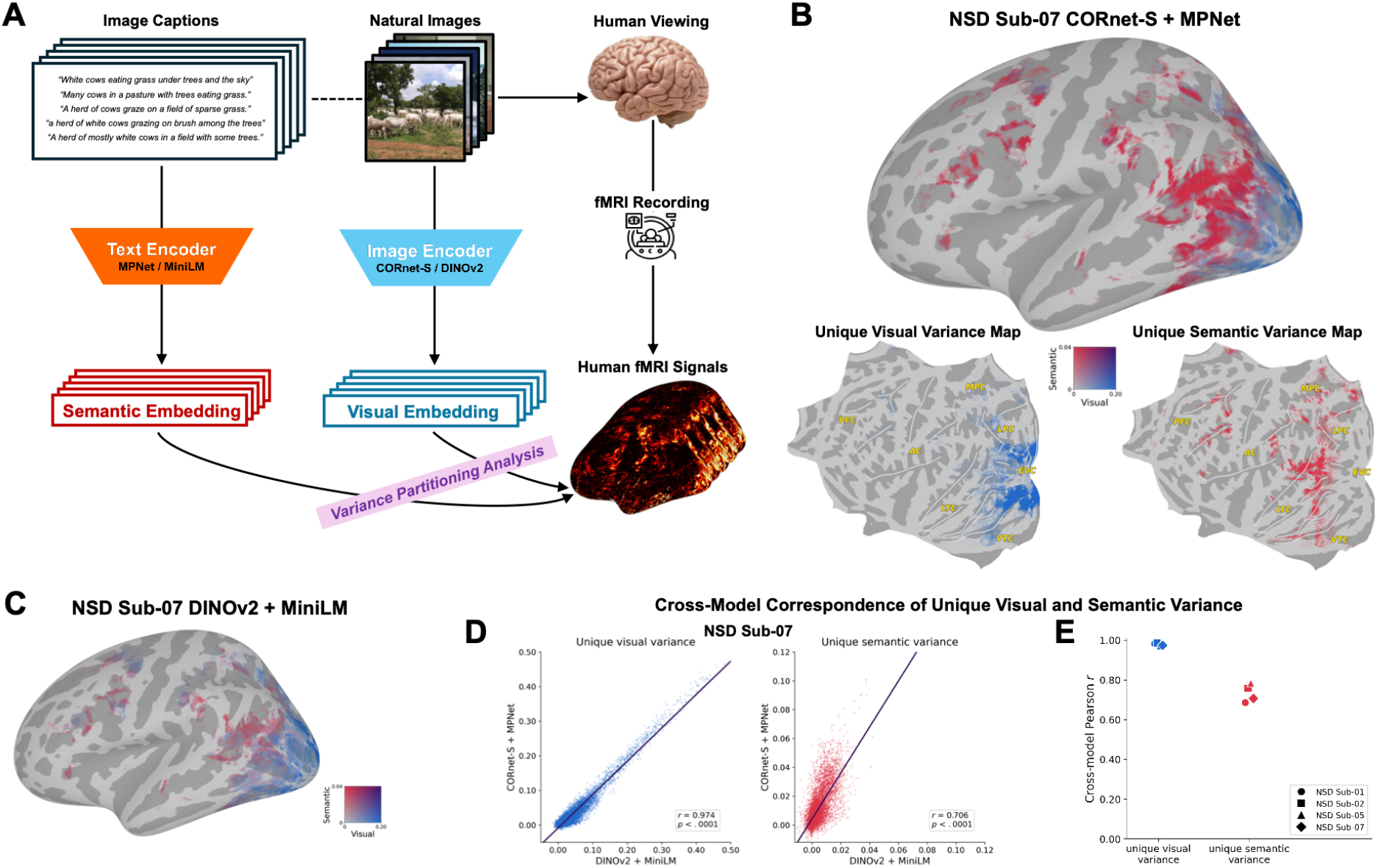
ANN-robust separation of visual and semantic representations during natural-scene viewing. (A) Image-to-fMRI encoding framework overview. Visual features were extracted from natural images using CORnet-S or DINOv2, whereas semantic features were extracted from image captions using MPNet or MiniLM. Separate visual, semantic, and joint encoding models predicted repeated-image-averaged 7T fMRI responses, and variance partitioning decomposed prediction performance into variance uniquely attributable to visual and semantic features. (B) Unique visual and unique semantic variance maps for NSD Sub-07 obtained with CORnet-S and MPNet. The bivariate surface map jointly represents the two variance components: blue indicates predominantly visual-specific variance, red indicates predominantly semantic-specific variance, and purple indicates their co-occurrence. Flattened maps show the two components separately. Only vertices within the participant- and model-specific encoding-significance mask are displayed. (C) Corresponding bivariate map obtained with DINOv2 and MiniLM. All surface visualizations show the left hemisphere. (D) Vertex-wise correspondence between the two encoding frameworks for unique visual variance and unique semantic vatriance in NSD Sub-07, evaluated within the intersection of their encoding-significance masks. Lines indicate least-squares fits; reported values are Pearson correlations. (E) Cross-framework Pearson correlations for all full participants. Color denotes variance component and marker shape denotes participant.

To ensure that the resulting cortical organization did not depend on a particular artificial neural network architecture, we repeated the complete analysis with two families of models: CORnet-S (11) + MPNet (12) and DINOv2 (13) + MiniLM (14). Both model families produced a posterior-to-anterior transition from predominantly visual to more semantic unique variance (Figure 1B-C). And the two encoding frameworks yielded highly consistent individual-level variance maps, with strong correlations across model pairs (Figure 1D-E). All subsequent analyses were restricted to participant- and model-specific cortical regions in which the joint encoding model showed significant positive held-out prediction. Downstream analyses were conditional on vertices with positive, FDR-significant held-out joint-model performance. These maps therefore provided two architecture-robust axes with which to test energetic organization, while retaining each participant’s own encoding-significance mask rather than constructing a group-average cortical map.

### Visual and semantic representations occupy distinct mitochondrial environments

We next asked whether these representational axes corresponded to regional mitochondrial organization. MitoD and MRC were projected to the cortical surface and related separately to visual- and semantic-specific variance. Because both cortical representations and mitochondrial properties vary along broad sensory-to-transmodal gradients, associations were estimated after adjustment for the principal cortical functional gradient (PG1) (15). Group inference used the four participant effects and 100,000 Moran spectral randomizations matched to each participant’s mask and preserving spatial autocorrelation (16,17).

The two representational axes showed markedly different mitochondrial profiles. In the CORnet-S + MPNet analysis, unique visual variance was negatively associated with MitoD (mean partial *r* = −0.110, 95% bootstrap interval −0.141 to −0.078, spatial-FDR *q* = 0.016) and more strongly with MRC (*r* = −0.224, −0.247 to −0.204, *q* = 0.002). Both effects replicated with DINOv2 + MiniLM (MitoD: *r* = −0.083, −0.113 to −0.046, *q* = 0.043; MRC: *r* = −0.220, −0.249 to −0.188, *q* = 0.003; Figure 2A-B). Thus, cortical regions in which visual features explained more unique response variance tended to exhibit both lower mitochondrial abundance and lower mitochondria-specific respiratory capacity.

**Figure 2.**
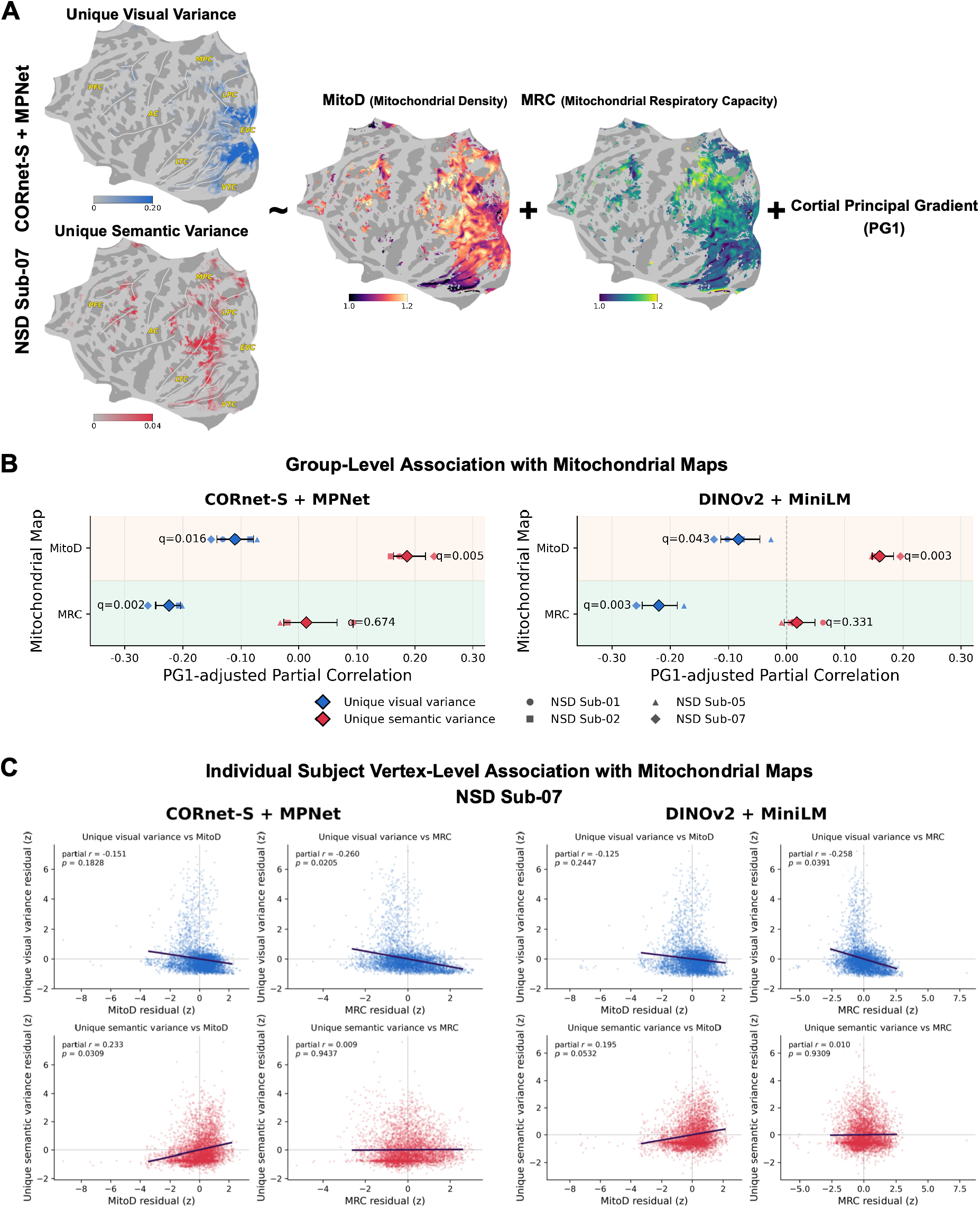
Distinct mitochondrial phenotypes align with visual- and semantic-specific cortical variance. (A) Analysis framework illustrated for NSD Sub-07 under the CORnet-S + MPNet encoding model. Unique visual and unique semantic variance maps were compared with MitoD and MRC while adjusting for the PG1. MitoD and MRC are displayed within the individual-specific encoding-significance mask. (B) Group-level PG1-adjusted partial correlations between each variance component and mitochondrial phenotype for CORnet-S+MPNet and DINOvs + MiniLM (*n* = 4 NSD participants). Small symbols show participant-level effects; symbol shape identifies participant. Large diamonds show inverse-Fisher-transformed group means, and error bars show 95% participant-bootstrap intervals. Spatial significance was evaluated using 100,000 mask-matched Moran spectral randomizations preserving cortical spatial autocorrelation. Displayed *q* values are two-sided spatial p values corrected by the Benjamini-Hochberg procedure across the prespecified family of four tests – two variance components by two mitochondrial phenotypes – separately within each encoding framework. (C) NSD Sub-07 residual-residual plots after PG1 adjustment, shown separately for the two encoding frameworks. Each point represents a cortical vertex. Lines indicate least-squares fits; annotations report partial Pearson correlations and two-sided participant-level spatial-surrogate *p* values.

In contrast, unique semantic variance showed a different mitochondrial profile. It was positively associated with MitoD in both model families (CORnet-S + MPNet: *r* = 0.186, 0.164 to 0.218, *q* = 0.005; DINOv2 + MiniLM: *r* = 0.160, 0.148 to 0.184, *q* = 0.003), whereas its association with MRC was near zero (*r* = 0.013 and 0.018; *q* = 0.674 and 0.331, respectively). Participant effects were directionally consistent (Figure 2C, an individual example of NSD Sub-07). Because MitoD and MRC were themselves moderately correlated within the analysis masks across NSD subjects (PG1-adjusted *r* = 0.568-0.666), these effects should not be interpreted as independent causal contributions of the two mitochondrial measures. Instead, they reveal distinct regional mitochondrial profiles associated with the cortical strength of visual versus semantic information representation.

The contrast between the two axes argues against a simple account in which stronger cortical representation is uniformly associated with greater energetic capacity. Visual-specific representation was strongest in regions with lower mitochondrial abundance and lower respiratory specialization, whereas semantic-specific representation preferentially tracked mitochondrial abundance without corresponding variation in MRC. Mitochondrial abundance and mitochondria-specific respiratory specialization therefore appear to capture separable dimensions of the biological organization associated with different forms of cortical information processing.

### Distinct transcriptional programs accompany the two representational axes

Building on the MitoMap analysis, we next turned to cortical gene expression to further probe the molecular basis of visual and semantic information encoding and to ask whether their cortical dissociation is reflected in transcriptomic organization. Microarray data from all six Allen Human Brain Atlas donors were processed with a donor-consistent workflow (18) and matched to each participant’s left-hemisphere encoding mask. Depending on participant and model, 169-248 samples from all six donors were retained (Figure 3A). For each of 16,008 quality-controlled genes, we estimated donor- and PG1-adjusted associations with unique visual variance or unique semantic variance separately in each participant, then ranked genes by the four-participant group *t* statistic. We used the complete rankings for competitive enrichment rather than selecting genes by an arbitrary significance threshold.

**Figure 3.**
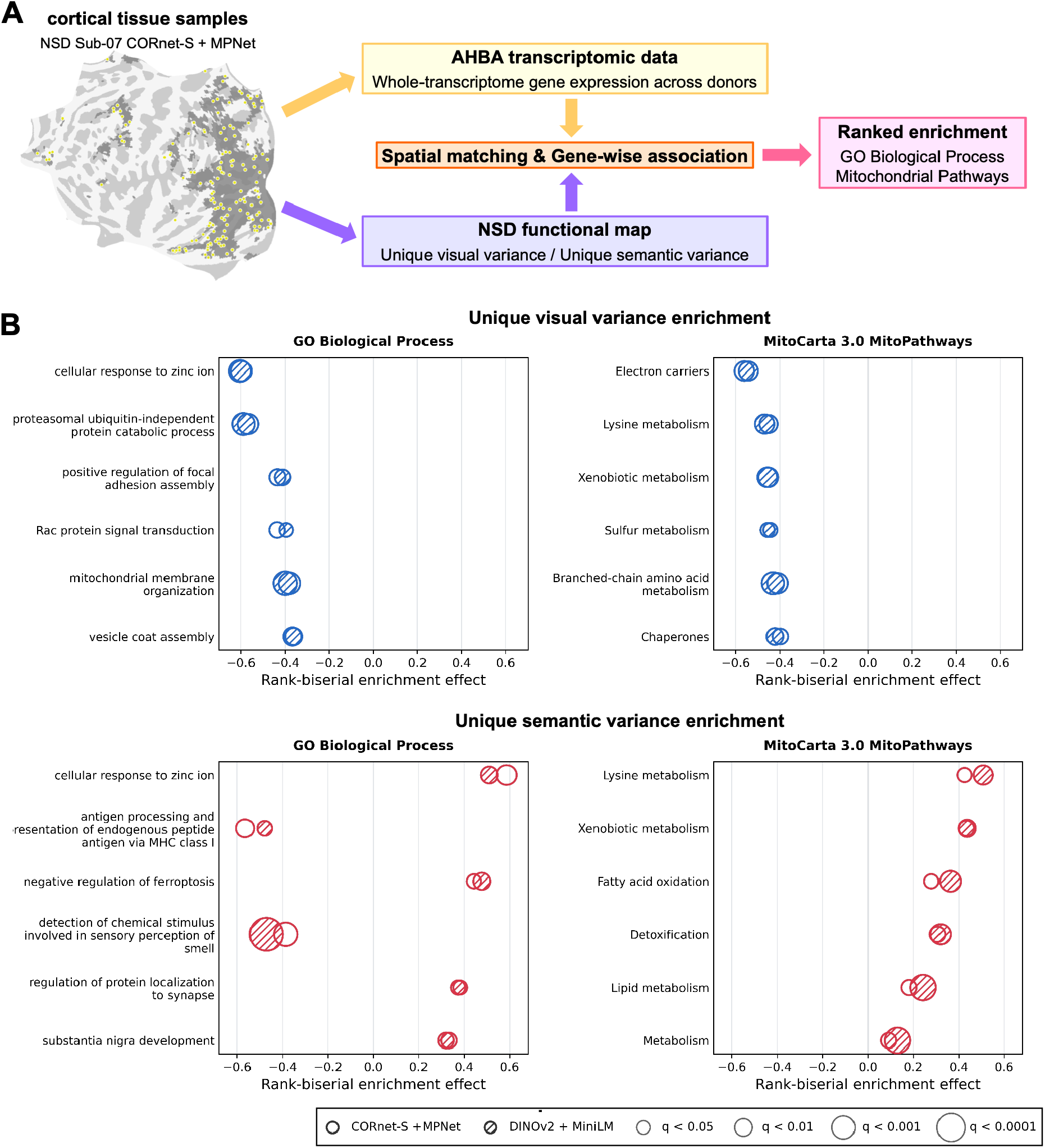
Transcriptomic programs associated with visual and semantic representational axes. (A) Transcriptomic analysis workflow. Yellow points show the 169 AHBA cortical tissue samples from all six donors that were spatially matched to the left-hemisphere encoding-significance mask of NSD Sub-07 under the CORnet-S + MPNet framework. For each participant and encoding framework, retained AHBA samples were matched to the corresponding unique visual and unique semantic variance maps. Gene expression and variance values were adjusted for donor identity and PG1 before gene-wise association analysis. Participant-level associations were Fisher-transformed and summarized by a four-participant group *t* statistic, which ranked 16,008 genes for enrichment analysis. (B) Replicated enrichment of GO Biological Process terms and MitoCarta 3.0 MitoPathways for unique visual variance (blue) and unique semantic variance (red). The horizontal axis shows the rank-biserial effect from a competitive two-sided Mann-Whitney test; negative values indicate that genes in a set were shifted toward negative gene-variance associations, whereas positive values indicate that a shift toward positive association. Hollow circles denote the results based on CORnet-S + MPNet encoding framework, and hatched circles denote the results based on DINOv2 + MiniLM encoding framework. Bubble area represents −log10(*q*). Terms were required to reach *q* < .05 in both encoding frameworks and to show concordant effect directions; the six terms with the largest mean absolute rank-biserial effect across frameworks are displayed for each library and variance component. *q* values were obtained by Benjamini-Hochberg correction separately within each gene-set library, encoding framework, and variance component.

Gene-set results reinforced the phenotypic dissociation (Figure 3B), revealing different highly reproducible molecular axes underlying unique visual and semantic representations. Among terms significant in both model families with the same direction, Unique visual variance showed negative enrichment for mitochondrial membrane organization and for MitoCarta (19) pathways involving electron carriers, chaperones, branched-chain amino-acid, lysine, sulfur and xenobiotic metabolism. Unique semantic variance instead showed positive enrichment for MitoCarta pathways involving fatty-acid oxidation, lipid metabolism, detoxification, lysine and xenobiotic metabolism. GO terms (20) also differentiated the axes: Unique visual variance rankings emphasized protein catabolism, vesicle coats and focal-adhesion-related processes, whereas unique semantic variance rankings included positive ferroptosis regulation and synaptic protein localization, alongside negative antigen-presentation and olfactory-detection terms. These annotations are best read as pathway-level hypotheses rather than evidence that any single pathway mediates a representation.

## Discussion

Together, these findings extend the study of brain energetics from anatomical variation and intrinsic network organization to the content of cortical representations. The visual and semantic variance maps describe not simple where activity occurs, but which type of information uniquely accounts for stimulus-evoked responses at each cortical location. Their reproducible associations with mitochondrial phenotypes across two encoding frameworks suggest that the molecular organization of cortex is related not only to its anatomy and macroscale connectivity, but also to the information it preferentially represents during cognition. The accompanying transcriptomic results extend this correspondence to coordinated gene program providing a second molecular level at which the visual-semantic distinction is expressed.

The mitochondrial associations did not follow a simple “more energy, stronger representation” model. Visual-specific variance was negatively associated with both mitochondrial abundance and mitochondrial-specific respiratory capacity, whereas semantic-specific variance was positively associated with mitochondrial abundance but showed no reliable association with respiratory capacity. These contrasting profiles suggest that mitochondrial abundance and respiratory specialization capture different aspects of the cortical organization supporting information processing. The transcriptomic enrichments were consistent with this distinction: visual and semantic gene rankings showed opposing enrichment across several mitochondrial metabolic pathways and broader cellular processes. These effects should be interpreted as differences in regional molecular organization rather than as measurements of the energy consumed while participants viewed images.

Several considerations define the boundary of these conclusions. The analyses establish cross-scale spatial correspondence, not a causal effect of mitochondrial biology on visual or semantic representation. MitoD and MRC were obtained from a postmortem mitochondrial atlas, and AHBA expression profiles were obtained from a separate set of donors; neither resource provides tissue measurements from the four NSD participants. The small number of intensively sampled participants supports within-participant mapping and cross-framework replication but could limit population-level generalization. PG1 adjustment and spatial-autocorrelation-preserving inference reduce confounding by broad cortical gradients, but they cannot eliminate unmeasured cytoarchitectural, vascular, or cellular factors shared across maps. The transcriptomic enrichments therefore identify pathway-level hypotheses rather than mechanisms. Establishing causality will require designs that measure mitochondrial phenotypes and information-selective neural responses in the same individuals, together with experimental manipulation or longitudinal assessment of mitochondrial function.

Despite these limitations, the convergence across precision functional mapping, two independent representational models, mitochondrial phenotypes, and genome-wide expression suggests that cortical specialization extends across levels of organization. Regions that preferentially represent visual versus semantic information are not distinguished only by their functional responses; they also occupy different energetic and molecular landscapes. These findings provide a bridge between systems-level descriptions of human cortical representation and the cellular architecture that supports neural computation.

## Materials and Methods

### Study design and inferential framework

We tested whether cortical variance specifically attributable to visual or semantic information during natural-scene viewing is spatially aligned with regional mitochondrial phenotypes and cortical transcriptional organization. The analysis proceeded in four stages. First, participant-specific visual-only, semantic-only and joint image-to-brain encoding models were fitted and used for variance partitioning. Second, downstream analyses were restricted to participant- and model-specific cortical masks defined by significant held-out performance of the joint encoding model. Third, the resulting variance maps were related to mitochondrial density (MitoD) and mitochondrial respiratory capacity (MRC) (8), with the principal cortical functional gradient (PG1) included as a covariate and spatially autocorrelation-preserving null models used for inference. Fourth, the same variance maps were related to genome-wide expression in the Allen Human Brain Atlas (AHBA) (9), followed by competitive enrichment analysis of complete gene rankings.

Two complementary visual–semantic feature combinations – CORnet-S + MPNet and DINOv2 + MiniLM – were carried through the full workflow. They were treated as parallel analyses rather than pooled as independent observations: cortical masks, participant-level effects, spatial nulls and false-discovery-rate (FDR) families were defined separately for each feature combination. Group-level inference was based on participant-level effects from the four intensively sampled Natural Scenes Dataset (NSD) (10) participants, not on correlations computed from a participant-averaged cortical map.

### NSD participants, task and MRI acquisition

We analyzed the four NSD participants who completed all 40 functional scanning sessions (Sub-01, Sub-02, Sub-05 and Sub-07). In the original NSD experiment, participants viewed color natural scenes drawn from Microsoft COCO while performing a continuous old–new recognition task. Images subtended 8.4° × 8.4°, were presented for 3 s and were separated by a 1-s interval. Each participant completed 750 trials per session, yielding 30,000 trials and approximately 10,000 distinct images; every retained image was presented three times. Functional MRI was acquired at 7T using whole-brain gradient-echo echo-planar imaging (1.8-mm isotropic voxels; repetition time, 1.6 s). Full acquisition, preprocessing, consent and ethical-approval procedures are described in the NSD data paper (10). The present secondary analysis used only publicly released, de-identified data.

### Single-image response estimates and train–test split

We used NSD single-trial beta estimates from the *betas_fithrf_GLMdenoise_RR* release in fsaverage surface space (163,842 vertices per hemisphere). These estimates incorporate voxel-wise hemodynamic-response-function fitting, GLMdenoise and ridge regression. Values were divided by 300 according to the NSD storage convention. Before any encoding analysis, the three beta estimates corresponding to the same image were averaged, so that each image contributed one response vector per participant and hemisphere.

The official set of 1,000 images viewed by all four participants (shared1000) was used as a fixed held-out test set. All other participant-specific, non-shared images not flagged by NSD constituted the training set. The resulting training sets contained 8,997, 8,995, 8,996 and 8,993 images for Sub-01, Sub-02, Sub-05 and Sub-07, respectively; all participants contributed the same 1,000 test images. Feature reduction, ridge-penalty selection and stacking-weight estimation used training images only. The held-out images were used for final prediction, variance partitioning and definition of the encoding-significance mask. The initial candidate cortex contained all non-medial-wall vertices (149,955 in the left hemisphere and 149,926 in the right hemisphere).

### Visual and semantic feature extraction

#### CORnet-S + MPNet analysis

Visual features were extracted with a pretrained CORnet-S (11). Images were resized to 256 pixels using bicubic interpolation, center-cropped to 224 × 224 pixels, converted to tensors and normalized using ImageNet channel means and standard deviations. Activations were captured from the V1, V2, V4 and IT modules and summarized with the same 1 × 1, 2 × 2 and 4 × 4 spatial-pyramid average pooling, producing four visual feature groups. Semantic features were extracted with MPNet (sentence-transformers/all-mpnet-base-v2) (12). Each NSD image was associated with five COCO captions. The five captions were embedded separately without length normalization, and their embeddings were arithmetically averaged to produce one 384-dimensional semantic vector per image.

#### DINOv2 + MiniLM analysis

The second analysis used a pretrained DINOv2 ViT-B/14 (13) for visual features. Image preprocessing matched the CORnet-S pipeline. Images were resized to 256 pixels using bicubic interpolation, center-cropped to 224 × 224 pixels, converted to tensors and normalized using ImageNet channel means and standard deviations. Patch-token representations were obtained from transformer blocks 3, 6, 9 and 12; the class token was excluded. Each reshaped two-dimensional activation map was summarized by adaptive average pooling over 1 × 1, 2 × 2 and 4 × 4 spatial-pyramid levels and concatenated, producing a separate visual feature group for each block. Semantic features were extracted with MiniLM (sentence-transformers/all-MiniLM-L6-v2) (14). Each NSD image was associated with five COCO captions. As above, the five non-length-normalized caption embeddings for each image were averaged. Features were generated once for the union of 36,981 images retained across the four participants and subsequently indexed into each participant’s training and test sets.

### Encoding models, nested cross-validation and variance partitioning

Encoding models were fitted separately for each participant, hemisphere and feature combination. Each visual feature group and the semantic feature group were reduced independently to at most 512 principal components using randomized singular-value decomposition with whitening. Principal-component analysis (PCA) was nested to prevent information leakage. The participant-specific training set was partitioned into five shuffled outer folds. For each fold, PCA was fitted only to the outer-training images and applied to the corresponding validation images. PCA for final test-set prediction was fitted to the complete training set and then applied to the 1,000 held-out images.

For every feature group and cortical vertex, we fitted ridge regression with an intercept. The ridge parameter was selected separately for each vertex from 0.01, 0.1, 1, 10, 100, 1,000, 10,000 using four-fold cross-validation within the relevant training partition, minimizing summed squared prediction error. The five outer folds generated out-of-fold predictions for every training image. A model refitted to all training images generated predictions for the held-out test set.

The out-of-fold predictions from the four visual feature groups were combined vertex-wise by convex stacking. Stacking coefficients were constrained to be non-negative and sum to one, and a free intercept was fitted. The single semantic prediction was calibrated using the same out-of-fold procedure. A joint model combined all four visual predictions and the semantic prediction in one convex stack. Thus, for every cortical vertex, we obtained visual-only, semantic-only and joint predictions without using test-set responses for model fitting or hyperparameter selection.

Held-out predictive performance was quantified as

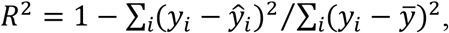

where *y*_*i*_ and *ŷ*_*i*_ are the measured and predicted responses to test image *i*. Held-out Pearson prediction correlations were retained for quality control, whereas variance partitioning and mask definition used predictive *R*^2^. Vertex-wise components were

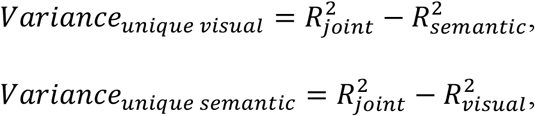

and

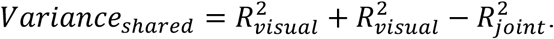

Unique visual variance and unique semantic variance therefore quantify held-out cortical variance uniquely attributable to the visual and semantic feature spaces, respectively. Shared denotes the overlap assigned to both feature spaces by commonality analysis. Raw component estimates, including negative values, were retained rather than clipped. The principal manuscript analyses focus on unique visual variance and unique semantic variance; Shared was nevertheless computed throughout and retained in the predefined multiple-testing families so that multiplicity correction did not become less stringent after focusing the presentation.

### Participant- and model-specific encoding-significance masks

Downstream molecular analyses were restricted to cortex reliably predicted by the joint encoding model. For each participant, feature combination and non-medial-wall vertex, we generated 10,000 paired bootstrap resamples of the 1,000 test images. Each resample drew 1,000 images with replacement and applied the same resampling counts to measured responses and joint-model predictions before recomputing predictive *R*^2^. The one-sided empirical probability was

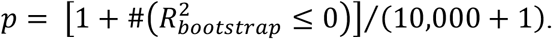

The left- and right-hemisphere candidate vertices were concatenated, and Benjamini– Hochberg (BH) FDR correction was applied separately for every participant and feature combination. Vertices with *q* < .05 and observed *R*^2^ > 0 formed the bilateral encoding-significance mask. Mask sizes ranged from 82,054 to 111,544 vertices across participants and feature combinations. Mask definition depended only on held-out joint-model prediction and not on unique visual variance, unique semantic variance, mitochondrial maps or gene expression. Because masks differed across participants and feature combinations, all downstream associations were estimated within the relevant mask and combined only at the participant-effect level.

### Cross-model correspondence of variance maps

To quantify the robustness of the functional decomposition to the chosen feature architectures, we compared the CORnet-S + MPNet and DINOv2 + MiniLM variance maps within each participant. Unique visual variance values from the two analyses were correlated across the bilateral intersection of their encoding-significance masks; unique semantic variance values were analyzed analogously. Pearson correlations and their conventional two-sided probabilities were calculated for descriptive purposes. Because cortical vertices are spatially autocorrelated, these conventional probabilities were not used as spatially corrected inferential evidence; robustness was assessed primarily from the magnitude and consistency of correlations across the four participants.

### Mitochondrial maps and cortical-hierarchy covariate

Mitochondrial maps were obtained from the MitoBrainMap (8). The source study assayed mitochondrial phenotypes in 3 × 3 × 3-mm tissue voxels from a frozen human coronal hemisphere section and used multimodal neuroimaging predictors to extend the measured phenotypes to brain-wide MNI152 maps. We analyzed two complementary atlas-derived phenotypes. MitoD is a composite measure of regional mitochondrial abundance derived from mitochondrial-density indicators. Tissue respiratory capacity (TRC) summarizes oxidative-phosphorylation (OXPHOS) enzyme activity per unit tissue, and MRC is the derived ratio MRC=TRC/MitoD. In biological terms, MRC was interpreted by the source study as mitochondria-specific specialization for OXPHOS energy transformation on a per-mitochondrion basis.

The MitoD and MRC brain-wide NIfTI maps were linearly projected from MNI152 volume space to the fsaverage10k cortical surface using neuromaps (21). For each map, a binary volume indicating non-zero source support was projected in parallel. Vertices with interpolated support ≤ 0.5 were excluded, and retained projected values were divided by the support weights to reduce attenuation at map boundaries.

PG1, the principal resting-state functional-connectivity gradient (15), was used as a covariate to account for the dominant unimodal-to-transmodal cortical hierarchy. PG1 was transformed from fsLR-32k to fsaverage10k using linear interpolation. Participant variance maps were transformed from fsaverage164k to fsaverage10k using linear resampling with correction for cortical-support weights; binary encoding masks were transformed by nearest-neighbor interpolation. The final vertex set for each participant and feature combination required encoding significance, non-medial-wall support and valid MitoD, MRC and PG1 values. This yielded 5,045–6,874 bilateral fsaverage10k vertices per participant and feature combination.

MitoD and MRC were analyzed in separate models. Accordingly, each effect estimates the association of a variance map with one mitochondrial phenotype after adjustment for PG1, not the association of that phenotype after conditioning on the other mitochondrial measure. This choice avoids assigning unstable “independent” effects to correlated, biologically related atlas phenotypes.

### Associations between variance maps and mitochondrial phenotypes

For each participant, feature combination, variance component and mitochondrial map, we calculated the Pearson partial correlation between the variance values and the mitochondrial values after linearly residualizing both variables with respect to an intercept and PG1. Unadjusted Pearson correlations were retained as descriptive results. The primary family contained unique visual variance and unique semantic variance crossed with MitoD and MRC (four tests) within each feature combination.

Group inference was performed on the four participant-level correlations. Correlations were Fisher z-transformed, and their consistency relative to zero was summarized with a one-sample t statistic (3 degrees of freedom). The group effect size was 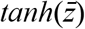. Ninety-five percent confidence intervals were obtained from 100,000 bootstrap resamples of participants with replacement. These intervals describe the stability of the mean participant effect and are necessarily coarse for n = 4.

Spatial significance was assessed with 100,000 Moran spectral randomizations (16,17). A separate null model was fitted to every participant- and feature-combination-specific mask. Within each hemisphere, pairwise geodesic distances were calculated on the fsaverage10k pial mesh and converted to inverse-distance weights; weights between hemispheres were set to zero. MitoD and MRC were randomized jointly with common Moran spectral sign flips, which preserved their spatial autocorrelation and mutual spatial relationship. For every permutation index, each participant received an independently generated, mask-matched mitochondrial surrogate. Partial correlations and the four-participant group t statistic were then recomputed. Two-sided add-one empirical probabilities were calculated as (*b*+1)/(100,000+1), where *b* is the number of null statistics at least as extreme in absolute value as the observed statistic. BH correction was applied across the four tests separately within each feature combination. No whole-cortex spin test, vertex-wise shuffle or participant sign-flip test was used.

### AHBA transcriptomic preprocessing

Genome-wide expression analyses used microarray data from all six AHBA donors. Because only two donors have substantial right-hemisphere sampling, analyses were restricted to left cortex. AHBA samples were assigned corrected MNI coordinates with abagen (22), anatomically mismatched samples were removed, and only samples annotated as left cortical tissue were retained.

Probe processing was independent of the fMRI results. Probes were reannotated, and probes without a valid Entrez identifier were discarded. A probe was required to exceed the AHBA background threshold in at least 50% of all retained left-cortical samples. Where multiple probes indexed the same gene, a single common probe was selected by highest differential stability across donors. Within each donor, expression was first normalized across genes for each tissue sample and then across left-cortical samples for each gene using scaled robust sigmoid normalization. Genes with non-finite values or near-zero variance in any donor were removed. We did not apply an additional gene-level differential-stability percentile cutoff. The processing sequence reduced 58,692 raw probes to 45,821 reannotated probes with valid Entrez identifiers, 32,917 probes passing the background criterion and a final universe of 16,008 genes measured across 1,312 left-cortical samples.

### Matching AHBA samples to participant-specific functional maps

Corrected sample coordinates were mapped to the nearest left-hemisphere fsaverage10k vertex using the MNI152-to-fsaverage10k regfusion coordinates provided by neuromaps (21). A sample was retained when its Euclidean distance to the nearest surface coordinate was ≤ 3 mm and that vertex fell inside the relevant participant- and feature-combination-specific encoding-significance mask. Unique visual variance, unique semantic variance, and PG1 were sampled at the matched vertex. This procedure was repeated independently for every participant and feature combination; a common intersection mask across participants was not imposed.

The retained sample counts were 218, 209, 248 and 204, 205, 234 and 169 in the CORnet-S + MPNet analysis, and 177 for NSD Sub-01, Sub-02, Sub-05 and Sub-07 in the DINOv2 + MiniLM analysis. Every participant–model analysis retained samples from all six AHBA donors. Distinct tissue samples mapping to the same surface vertex remained separate expression observations, whereas they were assigned the same spatial-surrogate value during permutation.

### Genome-wide gene–representation associations

Gene associations were estimated separately for each participant, feature combination, variance map and gene. For a given participant–model mask, the functional values and each gene’s expression values were residualized with respect to an intercept, five donor indicator variables and PG1. Pearson correlation between the two residual vectors yielded the donor- and PG1-adjusted gene association. Donor indicators remove mean expression differences among postmortem brains; they do not imply that AHBA donors and NSD participants were individually matched.

For each gene, the four participant-level partial correlations were Fisher z-transformed and entered a one-sample t statistic against zero. The reported group effect was 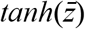. Confidence limits were obtained by enumerating all 4_4_ = 256 participant-level bootstrap resamples and taking the 2.5th and 97.5th percentiles. Group-level consistency, rather than a four-of-four conjunction rule, defined the primary transcriptomic result.

Gene-level spatial inference used a separate Moran spectral null for each participant and feature combination. Inverse geodesic-distance weights were constructed among the unique left-hemisphere fsaverage10k vertices linked to the retained AHBA samples. For every variance map, 100,000 spatial surrogates were generated. Samples linked to the same vertex shared the same surrogate value. At each iteration, donor- and PG1-adjusted gene correlations were recalculated for every participant, and the four-participant group t statistic was recomputed. Two-sided add-one empirical probabilities were obtained at both participant and group levels. Group probabilities were BH-corrected across all 16,008 genes separately for each feature-combination × variance-map panel. No gene was selected on the basis of its association before this genome-wide correction.

### Ranked gene-set enrichment analysis

Gene-set enrichment used the complete 16,008-gene ranking rather than a thresholded list of significant genes. Within each feature-combination × variance-map panel, genes were ranked by the signed four-participant group t statistic from the donor- and PG1-adjusted analysis. Three libraries were evaluated: Gene Ontology Biological Process (GO BP) (20),17 adult human cortical cell-type markers from the middle temporal gyrus taxonomy and MitoCarta 3.0 MitoPathways (19).

Human GO annotations were propagated through is a and part of parent relationships. Cell-type sets were formed from the union of single markers vs all, level4_markers_vs_all and combo_markers_vs_all markers and aggregated at Hodge hierarchy levels 1–3; identical marker sets were retained once using the most specific available label. Mitochondrial sets followed the MitoCarta 3.0 MitoPathways definitions. The global all-MitoCarta collection was excluded because it exceeded the common upper size limit. Across all libraries, only sets containing 10–500 genes mapped to the 16,008-gene background were retained, yielding 4,949 GO BP sets, 13 adult cortical cell-type sets and 72 MitoPathways.

For each set, a two-sided competitive Mann–Whitney test compared the ranks of member genes with the ranks of all other genes in the background. Rank-biserial correlation was used as the effect size: positive values indicate that set members are shifted toward genes positively associated with the variance map, whereas negative values indicate a shift toward the negative end. BH correction was performed separately within each library, feature combination and variance map. The enrichment test operates on gene ranks and does not treat AHBA tissue samples or cortical vertices as independent observations; spatial dependence at the cortical-sampling level was addressed in the preceding gene-association stage.

For presentation of GO BP results, redundant significant terms were greedily grouped separately for unique visual variance and unique semantic variance and separately by enrichment direction. Terms were grouped when the Jaccard similarity of mapped genes was at least .25 or the overlap coefficient was at least .5; the most statistically supported term was retained as the representative. This was a display-level reduction only and did not alter enrichment statistics or FDR values. For the main enrichment figure, a term additionally had to reach *q* < .05 in both feature combinations with concordant effect direction. Replicated terms were ranked by the absolute mean rank-biserial effect across the two feature combinations, and up to the six strongest GO BP representatives and six strongest MitoPathways were displayed for each variance axis. Cell-type and complete unfiltered library results were retained in the source tables.

### Statistical analysis, sample-size considerations and reproducibility

No statistical method was used to predetermine sample size. We included all four NSD participants who completed the full 40-session protocol, maximizing within-participant data while retaining a commonly held-out image set. No participant satisfying this criterion was excluded. The secondary analyses involved no experimental allocation, randomization or blinding. The encoding-mask bootstrap used a one-sided test because it tested whether held-out predictive *R*^2^ exceeded zero; all molecular association and enrichment tests were two-sided. Unless stated otherwise, significance was assessed at BH-FDR *q* < .05. Fixed random seeds were used for feature reduction, cross-validation, bootstrapping and spatial randomization. Numerical equivalence of the accelerated spectral-null implementation to explicit Moran-map reconstruction was verified before the 100,000-permutation analysis.

## Completing Interests Statement

The authors declare that no competing interests exist.

## Data Availability Statement

The processed data and code for this study is available at GitHub: https://github.com/ZitongLu1996/Mito_VisualSemanticRep.

